# Convergent temperature representations in artificial and biological neural networks

**DOI:** 10.1101/390435

**Authors:** Martin Haesemeyer, Alexander F Schier, Florian Engert

## Abstract

While discoveries in biological neural networks (BNN) shaped artificial neural networks (ANN) it is unclear if representations and algorithms are shared between ANNs and BNNs performing similar tasks. Here, we designed and trained an ANN to perform heat gradient navigation and found striking similarities in computation and heat representation to a known zebrafish BNN. This included shared ON and OFF type representations of absolute temperature and rates of change. Importantly, ANN function critically relied on zebrafish like units. We could furthermore use the accessibility of the ANN to discover a new temperature responsive cell type in the zebrafish cerebellum. Finally, our approach generalized since training the same ANN constrained by the *C. elegans* motor repertoire resulted in distinct neural representations matching features observed in the worm. Together, these results emphasize convergence of ANNs and BNNs on canonical representations and that ANNs form a powerful tool to understand their biological counterparts.

## Introduction

Neural network models such as the perceptron (Rosenblatt, 1962) and other parallel distributed processing models (Rumelhart et al., 1987) have been used to derive potential algorithmic implementations of cognitive processes, such as word perception (McClelland and Rumelhart, 1981; Rumelhart and McClelland, 1982) or attentional control (Cohen et al., 1990), using a neuronlike implementation. Importantly, these models demonstrated how complex computation could emerge from interconnected networks of simple units. This work strongly suggests that cognitive processes could be implicitly realized in trained connectivity weights rather than relying on a large diversity of computational units. Indeed, artificial neural networks (ANN) are increasingly successful in solving tasks long considered hallmarks of cognition in Biological Neural Networks (BNN). This includes visual discrimination tasks, playing chess and Go as well as spatial navigation (Silver et al., 2016; Krizhevsky et al., 2012; Trullier et al., 1997; Moser et al., 2008; Logothetis and Sheinberg, 1996; Banino et al., 2018; Cueva and Wei, 2018). While design principles of ANNs have been inspired by discoveries in biological neural networks (Hassabis et al., 2017), it is controversial whether both network types utilize the same fundamental principles and hence to what extent ANNs can serve as models of animal cognition (Lake et al., 2017). However, if representations and algorithms are shared between BNNs and ANNs, then artificial neural network models can be used to guide the analysis of large scale biological datasets that are becoming more and more prevalent in modern neuroscience (Engert, 2014). We recently used whole brain functional calcium imaging and circuit modeling to characterize how larval zebrafish detect and process temperature information to generate adaptive behavioral output (Haesemeyer et al., 2018). This approach uncovered critical neural temperature response type in the larval zebrafish hindbrain. These types broadly fall into two classes, ON and OFF cells that represent changes with opposing sign. Within each of these classes a set of neurons reports absolute temperature levels while another set encodes the rate of change. Since larval zebrafish readily navigate temperature gradients (Gau et al., 2013; Haesemeyer et al., 2015), we now generated and trained a deep convolutional neural network to solve a heat gradient navigation task using the behavioral repertoire of larval zebrafish. This approach allowed us to compare processing in biological and artificial neural networks that solve the same behavioral task. We found that these behavioral constraints led to striking similarities in temperature processing and representation in this ANN with the zebrafish biological circuits. Namely, the model parallels temperature representation in ON and OFF types as well as ANN units showing adapting and sustained responses effectively encoding both absolute temperature levels and rates of change. Importantly, ANN performance critically relied on units representing temperature in a fish-like manner while other nodes were dispensable for network function. We next used the accessibility of the ANN to uncover new features of the zebrafish BNN. This allowed us to identify a novel neuronal response type in the zebrafish brain that was predicted by the ANN but escaped detection in the previous brain wide calcium imaging experiments (Haesemeyer et al., 2018). Finally, our approach generalized since training the same ANN constrained by the *C. elegans* motor repertoire resulted in distinct neural representations that match closely features observed in the worm. On the one hand these results indicate that behavioral constraints can lead to convergence on canonical stimulus representations in ANNs and BNNs while they on the other hand demonstrate the utility of ANNs to gain insight into processing in biological networks.

## Results

### An ANN for heat gradient navigation

To compare temperature processing in ANNs and BNNs, we designed a branched, convolutional neural network processing sensory and behavioral history experienced over the past four seconds of exploring a temperature gradient (Figure 1a-d). We used rectifying linear units in the convolutional and hidden layers of the network to mimic neural rectification. The multi-layered feedforward structure of the network and usage of temporal convolution was inspired by general models of sensorimotor processing. We however did not match connectivity in the ANN to larval zebrafish circuitry to avoid constraining network representations by anatomy and to instead limit constraints to the goal of heat gradient navigation and the available motor repertoire. Previously, we observed heat responsive neurons that encode the direction of temperature change in the larval zebrafish hindbrain (Figure 1c, Fast ON and Fast OFF) which could be used for a simple form of prediction. We therefore designed our network to predict the temperature reached after enacting one of four possible zebrafish behavioral elements: Stay in place, swim straight, turn left or turn right. Importantly, this design choice is biologically plausible given the importance of behavioral forward models in decision making (Miall and Wolpert, 1996; Ahrens et al., 2012; Portugues and Engert, 2011; Mischiati et al., 2015) and allowed for supervised learning which greatly increased training efficiency compared to an approach directly reinforcing behavioral elements depending on navigation success.

**Figure 1:**
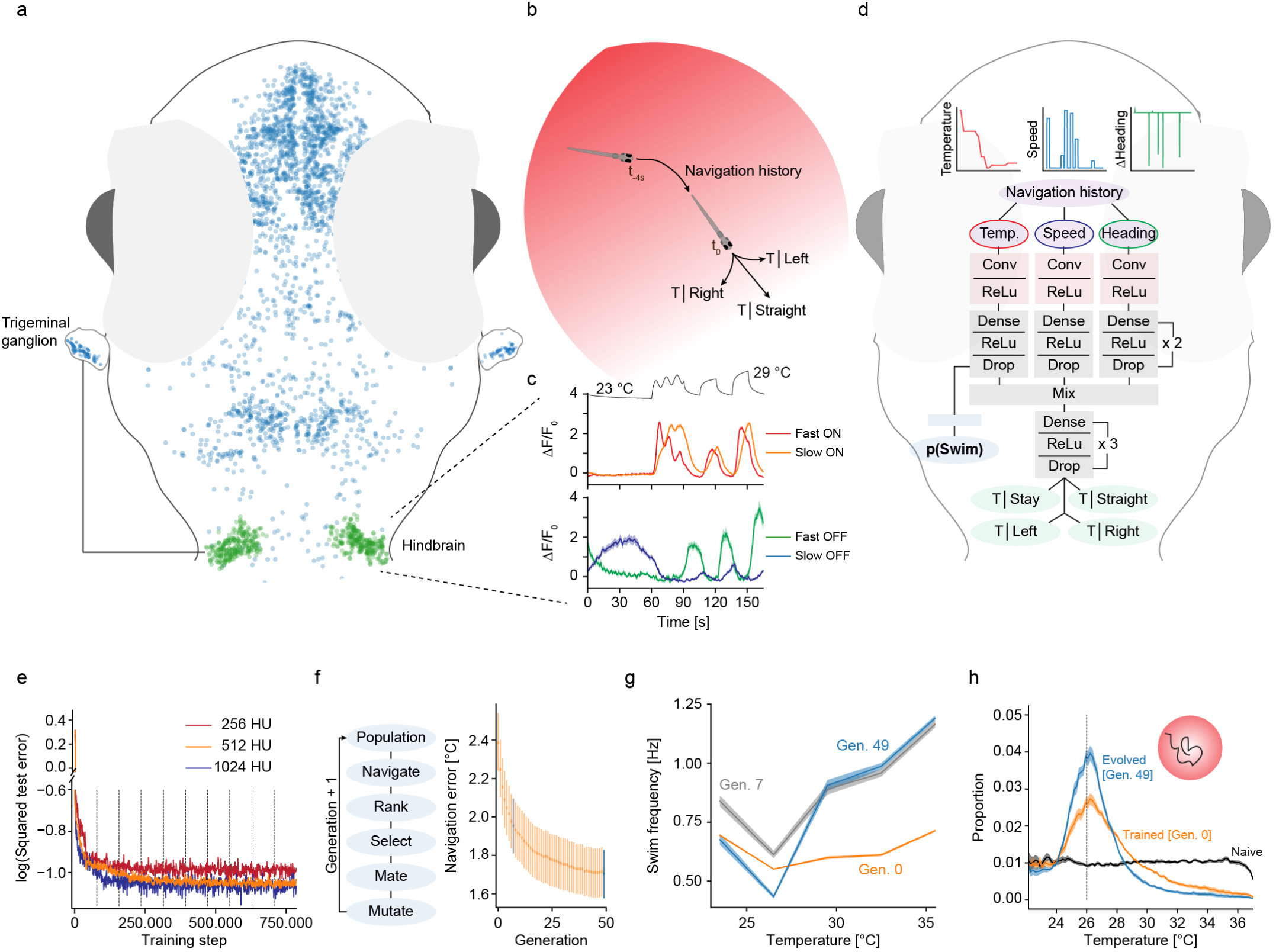
A deep convolutional network for gradient navigation a) Location of temperature modulated neurons (blue) in the zebrafish brain and sensory trigeminal ganglia. Temperature modulated neurons in a main processing area in the hindbrain are highlighted in green. b) Schematic of the task accomplished by the deep network. Given temperature and movement history in a heat gradient, the network predicts the resting temperature resulting from different behavior selections (stay, move straight, turn left, turn right). c) Zebrafish neuronal response types in the hindbrain region highlighted in a). Top panel: Adapting “Fast ON” (red) and nonadapting “Slow ON” (orange) neurons. Bottom panel: Adapting “Fast OFF” (green) and non-adapting “Slow OFF” (blue) neurons. Temperature stimulus presented to larval zebrafish depicted in black on top. d) Structure of the convolutional deep network for zebrafish temperature prediction. Curves on top depict example network input of the training dataset. Conv: Convolutional layer, ReLu indicates that network uses rectifying linear units, Drop: Indicates dropout used during training. e) Log of the squared error in temperature predictions (256 red, 512 orange, 1024 blue) on a test data set after the indicated number of training steps (dashed vertical lines demarcate training epochs). f) Evolutionary algorithm to learn weights that allow manipulation of bout frequency based on the output of the temperature branch (left panel) and progression of heat gradient navigation error as average distance from desired temperature across generations (right panel). Error-bars are bootstrap standard error across 20 evolved networks. Generation 7 highlighted in grey and last generation in blue for comparison with g) and h). g) For fully trained predictive network in generation 0 (orange), evolved generation 7 (grey) and network after completed evolution (blue) the average swim frequency produced by temperature in the gradient. Shading indicates bootstrap standard error across 20 networks. h) Occupancy in a radial heat gradient of naive (black), trained (orange) and evolved (blue) networks. Dashed vertical line at 26 *°*C indicates desired temperature. Shading indicates bootstrap standard error across 20 networks.

We trained the ANN using backpropagation on training data that was generated from a random walk through a radial heat gradient by drawing swims from distributions observed during realistic zebrafish heat gradient navigation (Haesemeyer et al., 2015). We used dropout (Srivastava et al., 2014) for regularization during training to mimic redundancy and noise generally observed in biological neural networks (BNNs). We compared training success in networks with different hidden layer sizes (256, 512 or 1024 units per layer), and since performance on a test dataset saturated with 512 hidden units per layer (Figure 1e), we chose this layer size for our ANN. Such networks successfully predicted the temperatures reached after typical swims with average errors *<* 0.1 *°*C. To transform this prediction into gradient navigation, we implemented a simple rule that favors those behaviors that bring the virtual fish closer to a set target temperature (Figure S1a). Invocation of this rule after training indeed led to efficient gradient navigation with an average distance from the target temperature of 2.4 *°*C compared to an average distance of 4.6 *°*C in naive networks.

While larval zebrafish swim in discrete bouts that occur, on average, at a frequency of 1 Hz, this baseline swim frequency is modulated by temperature (Haesemeyer et al., 2015), which could be a useful feature in the context of gradient navigation. This motivated us to extend the network by adding a module controlling swim probability independent of the already trained predictive function (Figure 1d, p(Swim)). To accomplish this task the p(Swim) module utilizes a set of weights that transform the output of the temperature processing branch of the network into a swim probability. In order to train this set of weights, we used an evolutionary algorithm since these can efficiently minimize empirical costs such as the heat gradient navigation error. This approach led to fast convergence in our task and after only 50 generations the algorithm converged on a specific set of weights for each of 20 trained networks (Figure 1f; Figure S1bd). Importantly, these weights led to increases in swim frequency as temperature departs from preferred values (Figure 1g), which is also observed in larval zebrafish (Gau et al., 2013; Haesemeyer et al., 2015). This significantly enhanced navigation performance of the network (Figure 1h) reducing the average distance from the preferred temperature from 2.4 *°*C to 1.7 *°*C. In summary, we designed and trained an artificial neural network to enable heat gradient navigation using a larval zebrafish behavioral repertoire.

### Processing and representation in the ANN parallels the zebrafish brain

After designing and training an ANN performing heat gradient navigation we sought to compare computation and stimulus representation within the ANN and the corresponding BNN. Previously, we characterized the computations underlying behavior generation during heat perception in larval zebrafish using white noise temperature stimuli (Haesemeyer et al., 2015). This approach allowed us to derive behavioral “filter kernels” that describe how larval zebrafish integrate temperature information to generate swim bouts (inset Figure 2a). These filter kernels revealed that larval zebrafish integrate temperature information over timescales of 500 ms to decide on the next swim and that they extract a derivative of the temperature stimulus as reflected in the 0-crossing of the filter (inset Figure 2a). The filter kernels furthermore indicated that swims were in part controlled by a strong OFF response just before the start of a bout. For comparison we now presented white noise temperature stimuli to the ANN and similarly computed filter kernels as behavior triggered averages for straight swims and turns (Figure 2a). These bear striking resemblances to the larval zebrafish filter kernels (Haesemeyer et al., 2015). Namely, even though no explicit constraints on integration timescales were given to the ANN, the behavior triggered averages reveal that most information is integrated over timescales less than a second, akin to larval zebrafish integration timescales (Figure S2b). This is likely the result of the ANN adapting to the average swim frequency of larval zebrafish used in the training data. Indeed, reducing the baseline swim frequency in the training data to 0.5 Hz elongates filter timescales, while an increase to 2 Hz heightens the filter peaks close to the swim start (Figure S2c). The ANN furthermore computes both a derivative and an integral of temperature and, notably, behavior is also influenced by a strong OFF response right before the start of a swim (arrowhead in Figure 2a). These are all hallmarks of larval zebrafish temperature processing. Furthermore as in zebrafish, the OFF response before the swim-start is more strongly modulated for turns than straight swims (Figure 2a), a strategy that zebrafish likely use to favor straight swims over turns when temperature approaches cooler, more favorable conditions (Haesemeyer et al., 2015). As expected, the swim triggered averages are completely unstructured in naive networks (Figure S2a).

**Figure 2:**
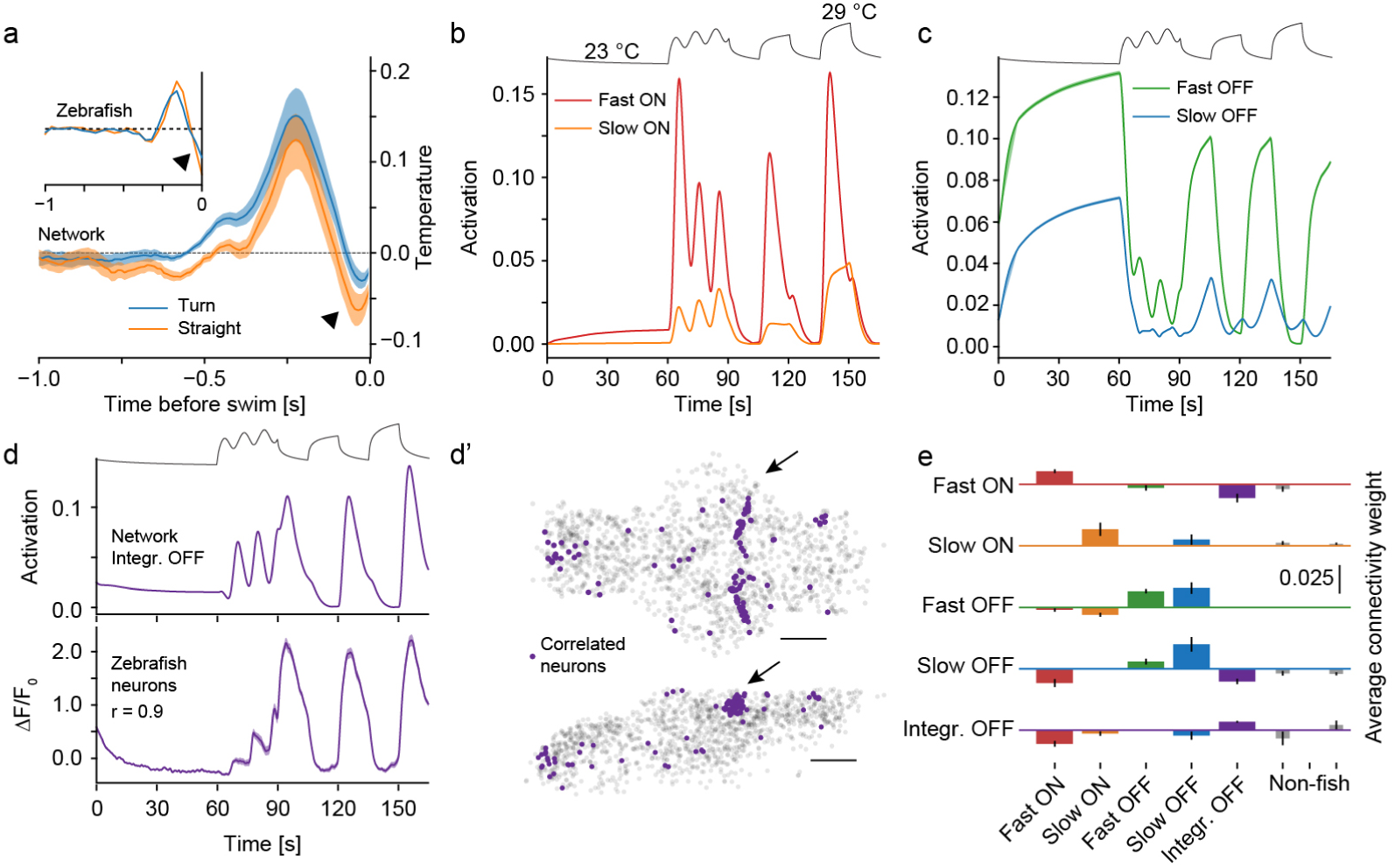
The network learns a zebrafish-like neural representation a) White noise analysis of behavior induced by the network depicting the average stimulus in the last second preceding a swim. Straight swim kernel orange, turn kernel blue. Inset shows zebrafish kernels for comparison with long straight bout kernel in blue and medium turn kernel in orange. Arrowhead indicates OFF response just before swim start in zebrafish and networks. b) Fish-like ON cell types revealed in the network when presenting the same temperature stimulus as presented to larval zebrafish (depicted on top for reference). Adapting Fast ON cells (red) and non-adapting Slow ON cells (orange). Compare with Figure 1c top panel. c) Fish-like OFF cell types revealed in the network. Adapting Fast OFF cells (green) and non-adapting Slow OFF cells (blue). Compare with Figure 1c bottom panel. d) Another cell type present in the network, “integrating OFF cells” (purple) can be used as a regressor (top panel) to identify the same, previously unidentified, cell type in zebrafish data (bottom panel, shading indicates bootstrap standard error across 146 zebrafish neurons). **d’)** The newly identified zebrafish cells cluster spatially, especially in a tight rostral band of the cerebellum (arrow). Top panel: Dorsal view of the brain (anterior left, left side bottom). Bottom panel: side view of the brain, anterior left, dorsal top). Scale bars: 100 *µ*m. e) Connectivity weights between layer 1 neuron types in the temperature branch (along x-axis) feeding into the indicated types of layer 2 neurons (panels). Fish-like types are indicated by corresponding colored bars and the three remaining non-fish like clusters are indicated by thinner gray bars on the right side. Errorbars indicate standard deviation. Shading indicates bootstrap standard error across 20 networks in all panels.

In the BNN of larval zebrafish we previously described a critical set of temperature encoding cells in the hindbrain consisting of ON and OFF type cells sensitive to absolute temperature levels on the one hand and changes in temperature on the other (Haesemeyer et al., 2018) (Figure 1c). Interestingly, spectral clustering across ANN units revealed a very similar representation in the temperature navigation ANN (Figure S2d-e). This included four prominent types. Two of which mimicked Fast ON and Slow ON activity found in the larval zebrafish hindbrain (Figure 2b) while another two paralleled Fast OFF and Slow OFF activity (Figure 2c). This similarity in stimulus encoding highlights convergence in representation and information processing between larval zebrafish and the designed ANN.

Encouraged by these similarities we tried to use other prominent response types found in the ANN to identify heat processing cells in the larval zebrafish brain that may have been missed by previous clustering approaches. In particular, the ANN contained two abundant response types that were quite different from cell types previously described in larval zebrafish: A group of ON-OFF units responding to both stimulus onand offset (Figure S2f) as well as a type that we termed “integrating OFF” as it was most active at low temperatures and integrated over successive temperature decreases (Figure 2d). We used the responses of these cell types as regressors to search the larval zebrafish brain data for cells with highly correlated responses. We couldnt identify cells that matched the response properties of the ON-OFF type since the most highly correlated cells in larval zebrafish rather resembled slow ON cells (Figure S2f). However, there was a group of cells with activity closely resembling the integrating OFF type (Figure 2d). Importantly, these cells clustered spatially in the larval zebrafish brain, where most of them were located in a tight band in the rostro-dorsal cerebellum (Figure 2d’, arrow). This anatomical clustering strongly supports the idea that these cells indeed form a bona-fide heat responsive cell type.

As in larval zebrafish we furthermore observed a clear asymmetry between encoding in ON and OFF type cells such that OFF cells were not the simple inverse of their ON cell counterparts (Figures 1c, 2b-c). Since our ANN used rectifying linear units which just like biological neurons cannot encode negative firing rates we wondered if this constraint caused this asymmetry. We therefore trained a set of networks in which we exchanged the activation function for the hyperbolic tangent function which results in an encoding that is symmetric around 0 (Figure S3a). These networks learned to predict temperatures and hence navigated heat gradients just as well as networks with rectifying linear units (Figure S3b-c) but remarkably they represented heat stimuli in a very different manner. Namely, OFF units in this network type were the exact mirror image of ON units (correlation *< -*0.99 for all pairs), which resulted in an overall simpler representation (Figure S3d-e). This notion of a simpler representation was supported by the fact that the first 4 principal components explained 99 % of the response variance across all cells in hyperbolic tangent networks while 7 principal components were needed in rectifying linear networks (Figure S3f). This suggests that the biological constraint of only transmitting positive neural responses shapes representations in ON and OFF type channels and increases required network complexity.

By design the connectivity of the ANN was not matched to the connectivity in the BNN of larval zebrafish, however analysis of connectivity weights between the hidden layers in the temperature branch of the network showed that zebrafish-like types receive on average stronger inputs from other zebrafish-like types than from non-fish types (Figure 2e). This suggests that zebrafish-like response types form a sub-network within the ANN.

The observed representations could be a general solution to navigational tasks that use time varying inputs or they could be specific to thermotaxis. To disambiguate these hypotheses we designed a network variant that has the exact same structure but a behavioral goal akin to phototaxis (Wolf et al., 2017; Huang et al., 2013; Chen and Engert, 2014). This network variant receives as input a history of angles to a lightsource and has the task of predicting the angular position after the same swim types used in the thermotaxis network (Figure S4a-b). We found that it efficiently learned to fixate a light-source and comparing cell responses between the thermotaxis and phototaxis networks revealed a much simpler stimulus representation in the latter (Figure S4c-e), arguing that such stimulus representations are not emergent features of networks trained to perform navigation but rather depend on the specific task at hand.

### Zebrafish like response types form the critical core of ANN function

After discovering clear parallels in representation and computation between ANNs and BNNs for thermal navigation we wanted to test the importance of the common response types for ANN function. Artificial neural networks are generally robust to random deletions of units and we expected this to specifically be the case in our ANNs as they were trained using dropout. Indeed, random removal of as much as 85 % of all units in the temperature processing branch had only a small influence on gradient navigation performance (Figure 3a). However, specific deletion of all Slow ON or Fast OFF like cells in the network, contrary to Fast ON, Slow OFF and Integrating OFF deletions, had a strong effect on temperature navigation (Figure 3b-c). Indeed, the Slow ON and Fast OFF types also have the highest predictive power on heat induced behaviors in the larval zebrafish hindbrain (Haesemeyer et al., 2018). Overall, deletion of any zebrafish like type in the network had a larger effect on gradient navigation performance than deleting individual types not found in larval zebrafish (Figure 3c) indicating a relatively higher importance of fish-like types. Strikingly, deleting all fish-like types in the temperature branch of the ANN nearly abolished gradient navigation while performance was hardly affected when deleting all non-fish types (Figure 3d). This demonstrates that fishlike response types are of critical importance for gradient navigation.

**Figure 3:**
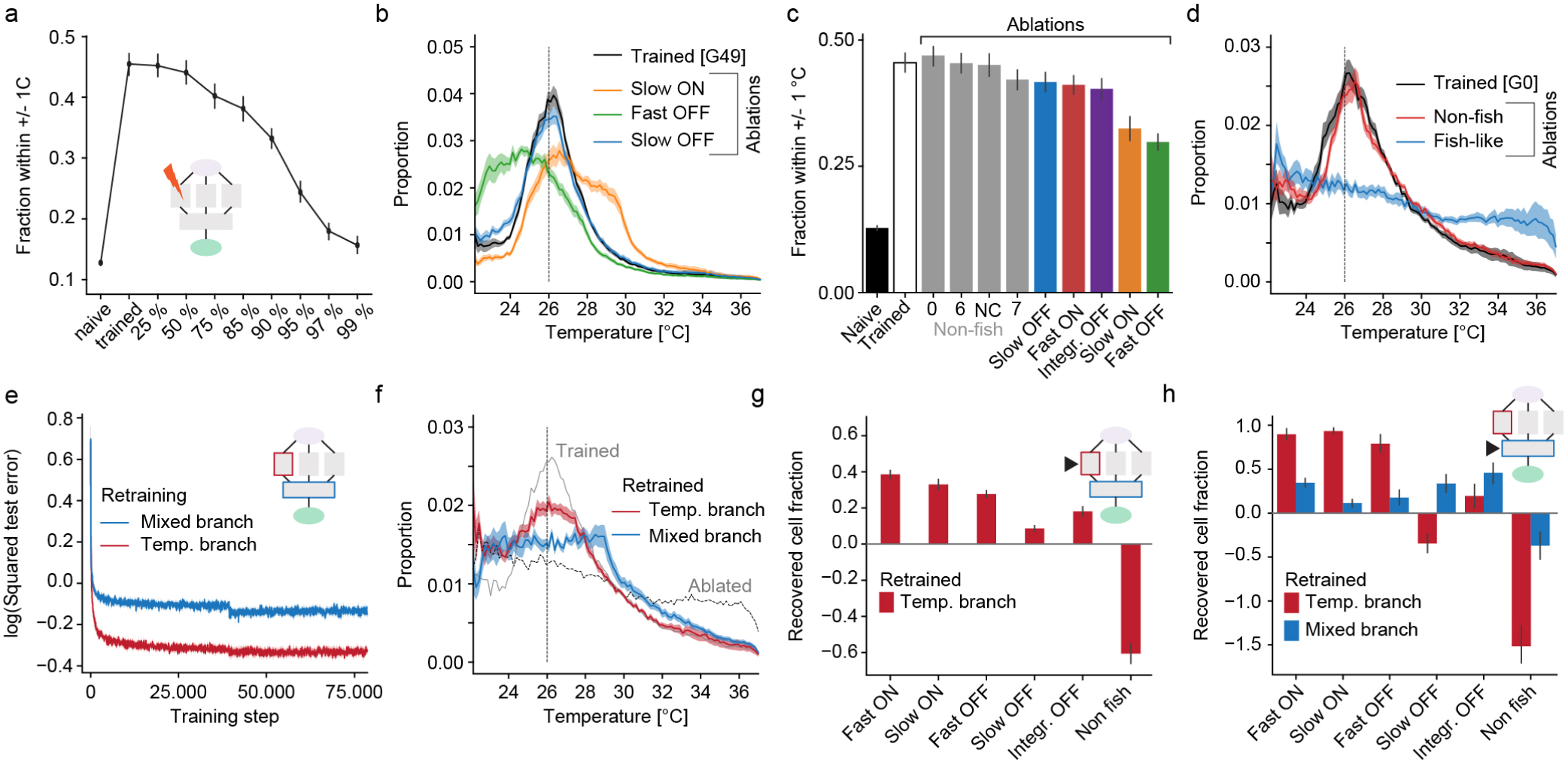
Ablations and retraining reveal importance of zebrafish like cell types a) Effect of random unit ablations on gradient navigation performance as fraction within 1 *°*C of desired temperature. Shown is performance for naive, fully trained and for random ablations of the indicated fraction of units in the temperature branch for zebrafish networks. Inset depicts location for all ablations. b) Occupancy in radial heat gradient for trained zebrafish networks (black) and after ablations of the indicated cell types (colored lines). c) Quantification of gradient navigation performance as fraction within 1 *°*C of desired temperature for naive and trained zebrafish networks as well as after ablations of the indicated cell types identified in larval zebrafish (colored bars) and those types not identified in fish (“Non-fish”), grey bars. Ablations are ordered according to severity of phenotype. d) Effect on gradient navigation of ablating all types identified in zebrafish (blue line) or all non-fish types (red line). Note that these are non-evolved networks to allow retraining analysis. Trained performance shown in black for reference. Since fish-like deletions would remove more units from the network overall than non-fish like deletions (63 vs. 29 % of all units) we matched the amount of ablated units in our non-fish ablations by additionally removing a random subset of units. e) Log of the squared error in temperature predictions of networks on the test data set after ablating all fish-like types in the temperature branch when either retraining weights in the temperature branch (red line) or in the mixed branch (blue line). Inset indicates retraining locations. f) Effect of re-training networks after ablating all zebrafish like neurons. Re-training was either limited to the temperature branch (red line) or the mixed branch (blue line). Solid grey line visualizes trained and dotted grey line ablated performance. **g-h)** Recovered fraction of individual cell types after retraining the temperature branch (red bars) or after retraining the mixed branch (blue bars). Insets depict retraining locations. g) Cell type fractions in temperature branch. h) Cell type fractions in mixed branch. Shading and error bars in all panels indicate bootstrap standard error across 20 networks.

To test whether the network could adjust to the absence of fish-like representations we performed localized retraining of the heat-navigation ANN, restricting updates to either the temperature branch of the network or the mixed branch that integrates temperature and movement information. One epoch of retraining improved network performance in both cases but retraining of the temperature branch led to considerably better prediction and gradient navigation performance compared with retraining of the mixed branch (Figure 3e-f). This difference indicates that while the temperature branch still transmits some usable information after the ablation of all fish-like types, the resulting representation of temperature is lacking information required for efficient navigation. To gain better insight into the consequences of retraining the network, we analyzed the distribution of response types in the temperature and mixed branch in the retrained networks. When retraining the temperature branch, fish-like types emerged at the expense of non-fish types giving further credence to their importance for temperature prediction and navigation (Figure 3g). Not surprisingly retraining of the temperature branch led to the reappearance of most fish-like types in the mixed branch as well (Figure 3h), again at the expense of non-fish types. Retraining the mixed branch however failed to re-generate most of the fish-like types indicating that these cannot be re-synthesized from information carried by non-fish types (Figure 3h). The only exceptions to this were Slow-OFF and Integrating-OFF cells which are the two cell types that receive fairly strong inputs from non-fish like types to begin with (Figure 2e).

### Generalization to a *C. elegans* network

To test whether our approach of behaviorally constraining an ANN generalizes to other species, we created a network variant using the behavioral repertoire displayed by *C. elegans* performing heat gradient navigation (Ryu and Samuel, 2002). The network had the same structure and task as the original network, predicting temperature after a movement (Figure 4 a-b) and was trained on a random walk through a heat gradient employing a *C. elegans* behavioral repertoire.

**Figure 4:**
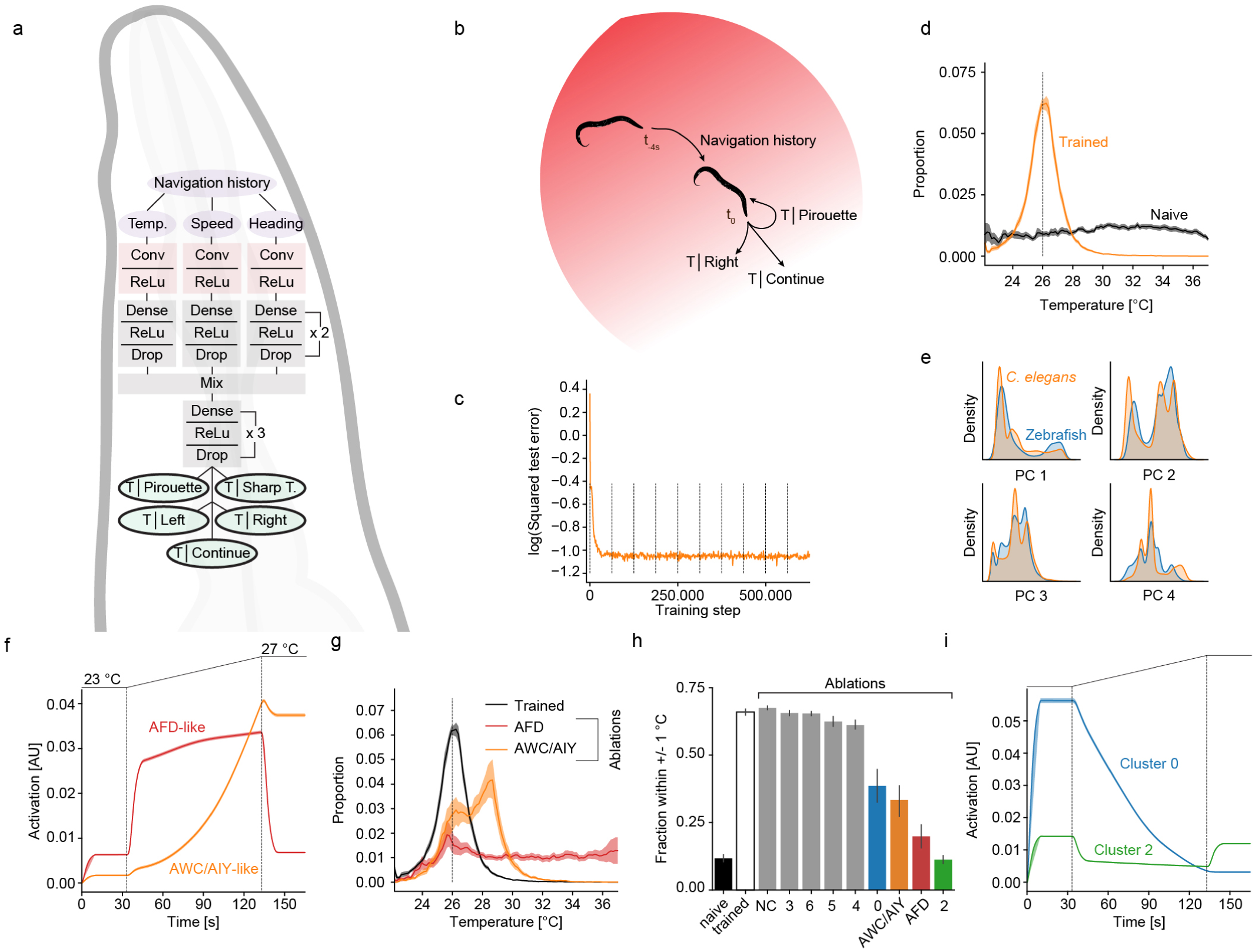
A network for *C. elegans* thermotaxis a) Architecture of the *C. elegans* deep convolutional network. Note that the architecture is the same as in Figure 1d except for the predictive output which is matched to the behavioral repertoire of *C. elegans*. b) Schematic of the task of the *C. elegans* ANN: The network uses a 4s history of experienced temperature and generated behaviors to predict the resting temperature after a *C. elegans* behavior (Continue moving, pirouette, sharp turn, small left turn, small right turn). c) Log squared error of temperature predictions on test data set during training. d) Occupancy in a radial heat gradient of naive (black) and trained (orange) *C. elegans* networks. Dashed vertical line at 26 *°*C indicates desired temperature. e) Comparison of all unit responses in the temperature branch of the zebrafish and *C. elegans* heat gradient ANN in PCA space when presenting the same time varying stimulus used in Figure 2b to both networks. The first four principal components capture *>* 95 % of the variance. Plots show occupational density along each PC for the zebrafish network (blue) and the *C. elegans* network (orange). f) Responses of two *C. elegans* like cell types when presenting a temperature ramp depicted in black on top. The red type shows adapting responses like the AFD neuron while the orange type reports temperature level as suggested for the AWC/AIY neurons. g) Occupancy in radial heat gradient for trained *C. elegans* networks (black) and after ablations of the indicated cell types (colored lines). h) Quantification of gradient navigation performance as fraction within 1 *°*C of desired temperature for naive and trained *C. elegans* networks as well as after ablations of the indicated cell types. Ablations are ordered by severity of phenotype. i) Responses of two *C. elegans* cell types the ablation of which results in strong gradient navigation phenotypes (k) to the same temperature ramp presented in f). Shading and error bars in all panels indicate bootstrap standard error across 20 networks.

Just like the zebrafish heat navigation ANN, the *C. elegans* ANN learned to predict temperatures when using a *C. elegans* behavioral repertoire (Figure 4c) and hence was able to navigate a heat gradient effectively (Figure 4d). Navigation performance was in fact better than for the zebrafish ANN (compare Figures 1h and 4d) which likely is a consequence of the expanded behavioral repertoire, especially the ability of trajectory reversals by executing pirouettes. We did not add an evolutionary algorithm to train changes in crawl frequency or speed since such behavioral modulation by temperature is not observed in the worm (Ryu and Samuel, 2002). Comparing responses in the temperature branches of zebrafish and *C. elegans* ANNs using principal component analysis revealed overlapping as well as divergent responses (Figure 4e). This partial overlap also became apparent when clustering cells from both networks and directly comparing the highest correlated clusters (Figure S5c-i). Here some clusters show near identical responses while other response types are exclusive to one of the two ANNs. Importantly, we could identify response types that represent temperature similarly to cells previously described in *C. elegans* (Figure S5a-b). This included a strongly adapting cell type that was most sensitive to changes in temperature similar to the *C. elegans* AFD neuron (Figure 4f) (Clark et al., 2006; Kimura et al., 2004).

Another cell type on the other hand largely reported absolute temperature levels as has been suggested for the AWC and AIY neurons (Figure 4f) (Kuhara et al., 2008). While the *C. elegans* ANN was as robust to random unit deletions as the zebrafish ANN (Figure S5k) it was considerably more sensitive to single cell type ablations. Removal of AFD like neurons severely reduced gradient navigation performance and especially affected cryophilic bias (Figure 4g), as reported for *C. elegans* itself (Chung et al., 2006). A weaker phenotype was observed when ablating AWC/AIY like neurons (Figure 4g-h) whose role in *C. elegans* thermotaxis is less well established (Garrity et al., 2010). The overall stronger dependence of network performance on individual cell types suggests a less distributed representation in the *C. elegans* ANN compared to the zebrafish ANN which was also mirrored in even sparser inter-type connectivity (Figure S5l). This may well be reflected in the animals themselves and in fact a recent paper applied a control theory paradigm to suggest links between *C. elegans* body bends and single motor neurons (Yan et al., 2017).

## Discussion

Artificial Neural Networks (ANN) and Biological Neural Networks (BNN) are very successful in solving problems of various complexity but how these networks accomplish such tasks is still largely unclear. Uncovering the fundamental principles that govern these operations is daunting in the case of BNNs because experimental access is limited and the underlying implementation is not necessarily aligned with human intuition. The principles underlying the operation of ANNs on the other hand are likely easier to dissect because they are made by man.

And indeed, models of interconnected neuron-like units have been used extensively to model cognitive processes (Rumelhart et al., 1987; Rosenblatt, 1962). These network models were generally constrained by the desired behavior of a cognitive system and used to study how cognition can emerge from networks of interconnected, simple units. This work revealed potential algorithms the brain might implement to perform behavioral tasks. However, it is unclear to what extent the responses of units in the models correspond to neuronal responses observed in biological brains; in other words to what extent biological brains and artificial neural networks converge on similar solutions of algorithmic implementation.

Recently, parallels between neural processing in the ventral visual stream and artificial networks for image classification have been discovered (Khaligh-Razavi and Kriegeskorte, 2014; Yamins and DiCarlo, 2016) and networks that learn a place-cell like encoding have been shown to give rise to entorhinal cell types such as grid cells (Banino et al., 2018). Here, we extended these approaches by constraining neural network models using a heat gradient navigation task and species specific behavioral repertoire and subsequently comparing processing in the ANN to a zebrafish whole brain imaging dataset.

Through this approach we could show that processing and representation in the thermotaxis ANN bears striking similarities to BNN representations in zebrafish and *C. elegans* depending on the available motor repertoire. This includes a clear parallel in stimulus representation between zebrafish hindbrain neurons and the zebrafish ANN on the one hand and response similarities between neurons known to be important for *C. elegans* heat gradient navigation and the corresponding ANN on the other hand. This strongly argues that stimulus representations in BNNs and ANNs converged on a canonical solution throughout evolution and training respectively and that stimulus representations are likely constrained by behavioral goals and motor repertoires (Rosenblatt, 1962).

Constraining an artificial neural network by a heat gradient navigation task specifically allowed us to form testable predictions about the larval zebrafish and *C. elegans* BNN. This led to the identification of a novel heat response type in the larval zebrafish cerebellum. At the same time the differential effects of deleting fish-like types on navigation performance allows for the generation of testable hypotheses about the relative importance of individual cell types in the zebrafish BNN especially since the two most important response types in the ANN (Slow ON and Fast OFF) are also most strongly implicated in temperature processing in larval zebrafish (Haesemeyer et al., 2018). Virtual ablations also make strong predictions about the role of two OFF cell types in thermal navigation of *C. elegans* (Figure 4i). OFF type cells have so far not been implicated in *C. elegans* thermal navigation but recent whole brain imaging data in response to thermal stimuli suggests that thermosensitive OFF types do exist in *C. elegans* (Kotera et al., 2016).

In summary, the strong parallel between ANNs and BNNs implies that artificial networks with their greater amenability to analysis and manipulation over BNNs can serve as powerful tools to derive neural principles underlying cognition in biological brains.

## Author Contributions

MH conceived the project in discussion with FE and AFS, and carried out all experiments, programming and data analysis. MH, AFS and FE interpreted the data and wrote the manuscript.

## Acknowledgements

MH was supported in part of this project by an EMBO Long Term Postdoctoral fellowship (ALTF 1056-10) and a postdoctoral fellowship by the Jane Coffin Childs Fund for Biomedical Research. Research was funded by NIH grants 1U19NS104653 to FE and 1DP1HD094764 to AFS. We thank Hanna Zwaka for the *C. elegans* drawing in Figure 4a. We thank Armin Bahl, Andrew Bolton, James Fitzgerald and Aravi Samuel for helpful discussions and critical comments on the manuscript.

## Materials & Methods

All neural networks were implemented in Tensorflow (Abadi et al., 2016) using Python 3.6. All data analysis was performed in Python 3.6 using numpy, scipy and scikit learn (Pedregosa et al., 2011) as well as matplotlib and seaborn for plotting.

### Behavior generation

All simulations were run at an update frequency of 100 Hz. Since zebrafish swim in discrete bouts, network predictions were only evaluated whenever swim bouts were instantiated. This occurred at a baseline frequency of 1 Hz (i.e. with a probability of 0.01 given the update frequency). Even though *C. elegans* moves continuously, behavioral modules are only selected with slow dynamics (Ryu and Samuel, 2002). Hence to reduce computational load, models and behavior selections were only evaluated with a frequency of 0.1 Hz (i.e. with a probability of 10^*-*4^ given the update frequency).

**Zebrafish** Zebrafish behavioral parameters were based on swim bouts enacted by freely swimming larval zebrafish during heat gradient navigation (Haesemeyer et al., 2015). When the selected behavior was “stay” the virtual fish stayed in place for one update cycle. For all other choices displacements were drawn at random in mm according to:

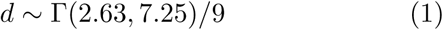

Turn angles (heading changes) for the three swim types were drawn at random in degrees according to:

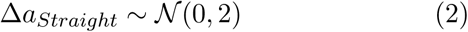

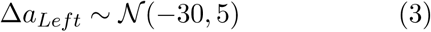

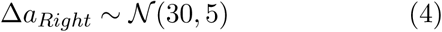

Each behavior was implemented such that each swim lasted 200 ms (20 timesteps). The heading change was implemented within the first frame while the displacement was evenly divided over all frames. This differs from true zebrafish behavior, where heading changes precede displacements as well but where both occur with distinct acceleration and deceleration phases.

The goal of these choices was to approximate larval zebrafish behavior rather than faithfully capture all different swim types.

***C. elegans*** *C. elegans* behavioral parameters were based on freely crawling worms navigating temperature gradients (Ryu and Samuel, 2002). When the selected behavior was “continue” or while no behavior was selected, the virtual worm was crawling on a straight line with heading jitter. The per-timestep displacement was drawn in mm according to:

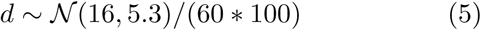

The per timestep heading jitter (random walk in heading direction space) was drawn in degrees according to:

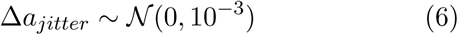

The other behaviors, pirouettes, sharp turns and shallow left or right turns were implemented as heading angle changes together with displacement drawn from the distribution above enacted over a total time of 1s. The heading angle changes were drawn in degrees at random as follows:

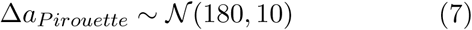

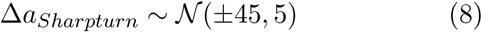

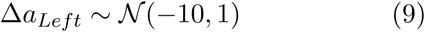

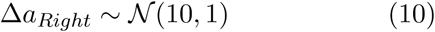

Again, the goal of these choices was to approximate *C. elegans* movement statistics during heat gradient navigation rather than faithfully recapitulating the full behavioral repertoire.

### Artificial neural networks

#### Structure

All networks had the same underlying architecture: Each input consisted of a 2D matrix, with 4 s of temperature, speed and delta-heading history at a simulation frame rate of 100 Hz. The first network operation was a mean-pooling, binning the inputs to a model frame rate of 5 Hz, the same frame rate at which larval zebrafish imaging data during heat stimulation was previously analyzed. After pooling, the inputs were split into the temperature, speed and delta-heading components and each component was fed into an identically constructed branch.

The input branches were designed with 40 linear rectifying convolutional layers each, that each learned one 1D filter over the whole 4s of history (20 weights). Convolution was performed such that only one output datapoint per filter (the full dot-product of the filter with the input) was obtained. These 40 outputs were subsequently passed through two hidden layers with 512 (or 256 or 1024) rectifying linear units. The outputs of all branches were subsequently concatenated and passed through another set of three hidden layers with 512 (or 256 or 1024) hidden units each before being fed into a linear output layer. This output layer consisted either of 4 linear units (for zebrafish networks) or 5 linear units (for *C. elegans* networks). The purpose of the output layer was to compute the temperature (or angle to a light source) 500 ms after enacting the chosen swim type in the case of zebrafish networks or 1 minute of straight continuous movement after enacting the chosen behavior in the case of *C. elegans* networks.

#### Training data generation and network training

For both zebrafish and *C. elegans* training data was generated by randomly choosing behavioral modules according to the statistics given above without any influence of temperature on behavioral choices. For training data generation the virtual animals explored two types of circular arenas with a radius of 100 mm each: In one arena, temperature increased linearly from 22 *°*C at the center to 37 *°*C at the periphery, while in the other arena the gradient was reversed. Training datasets were generated from the random walks through these arenas by simulating forward from each timestep for each possible behavioral choice. This way the true temperatures resulting from each behavioral choice were obtained together with the experienced temperature and behavioral history. For the zebrafish phototaxis ANN, the same strategy was employed but instead of recording temperature history and prediction, the angle to a light-source in the center of one arena with a radius of 100 mm was calculated. For each network type a test-data set was generated in the same manner to be able to evaluate prediction performance.

Networks were trained using stochastic gradient descent on mini-batches consisting of 32 random samples each. Notably, training was not successful when randomly mixing training data from both arena types. Every training epoch was therefore split into two halves during each of which only batches from one arena type training dataset were presented. We used an Adam optimizer [*learningrate* = 10^*-*4^, *β*_1_ = 0.9, *β*_2_ = 0.999, *ϵ* = 10^*-*8^] during training (Kingma and Ba, 2014), optimizing the squared loss between the network predicted temperature and the true temperature in the training dataset. Test batches were larger consisting of 128 samples each, drawn at random from both arena types.

Network weights were initialized such that gradient scales were kept constant across layers according to (Glorot and Bengio, 2010) by being drawn from a uniform distribution on the interval:

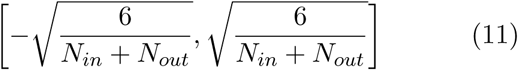

where *N*_*in*_ is the number of units in the previous and *N*_*out*_ the number of units in the current layer. For training regularization, we applied drop-out in all hidden layers, with a probability of 0.5 as well as weight decay, penalizing the squared sum of all weights in the network (*α* = 10^*-*4^). Networks were trained for 10 full epochs.

#### Navigation

The networks were used for heat gradient (or light) navigation in the following manner: For trained zebrafish networks each timestep had a probability of 0.01 of triggering a behavior (baseline movement frequency of 1 Hz after evolution this probability depended on the output of p(Move), see below). For *C. elegans* networks the probability of triggering a behavior was set to 10^*-*4^, resulting in a frequency of 0.1 Hz. If a timestep was not selected, zebrafish networks stayed in place while *C. elegans* networks continued to move as per the statistics above.

At each behavior evaluation, the preceding 4 s of sensory history as well as speed and delta-heading history were passed as inputs to the network. The network was subsequently used to predict the temperatures resulting from each possible movement choice (or the light angle in case of the phototaxis network). The goal temperature was set to be 26 *°*C and behaviors were ranked according to the absolute deviation of the predicted temperature from the goal temperature. For zebrafish networks, the behavior with the smallest deviation was chosen with a probability of 0.5, the 2^*nd*^ ranked with a probability of and the last two each with a probability of 0.125. For *C. elegans* networks, the highest ranked behavior was chosen with a probability of 0.5 the second highest with probability 0.2 and the remaining three behaviors with probability 0.1 each.

The chosen behavior was subsequently implemented according to the statistics above. Evaluations only resumed after a behavioral module was completed. Whenever a behavior would move a virtual animal outside the circular arena, the behavioral trajectory was reflected at the boundary.

#### Evolutionary algorithm to optimize control of swim frequency

A set of 512 weights was used to give the zebrafish networks control over swim frequency. A dot-product between the activations *a* of the last layer of the temperature branch of the network and these weights *w* was transformed by a sigmoid function to yield a swim frequency between 0.5 Hz and 2 Hz, by computing swim probabilities between 0.005 and 0.02 according to:

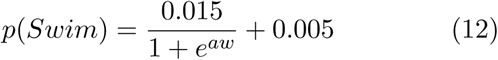

To learn a set of weights w that would optimize gradient navigation performance an evolutionary algorithm was used as follows:

1. Initialize 512 weight vectors, *w ∼𝒩* (0, 1)
2. For each weight vector run gradient simulation using it to control p(Swim)
3. Rank weight vectors according to average deviation from desired temperature
4. Pick 10 networks with lowest error and 6 networks at random
5. Form 16*16 mating pairs, mating each network with each other and with itself
6. For each mating pair generate two child vectors by randomly picking each vector element with probability 0.5 from either parent
7. Add random noise to each child vector, *ϵ ∼𝒩* (0, 0.1)
8. The 512 created child vectors form the next generation. Repeat from step 1.

Evolution was performed for 50 generations. The average across all 512 weight vectors in the final generation was used to control swim frequency during gradient navigation.

### Data analysis

For all thermotaxis network groups (Zebrafish ReLu and Tanh as well as *C. elegans*) a total of 20 networks were initialized and trained and all presented data is an average across these networks. For the phototaxis network a total of 14 networks was trained.

#### White noise analysis

For white noise analysis zebrafish networks were presented with randomly drawn temperature stimuli. As during navigation simulations networks enacted behaviors based on temperature prediction at a frequency controlled by p(Move). The stimulus used for white noise presentation was modeled after the stimulus used previously in freely swimming larval zebrafish (Haesemeyer et al., 2015), however since there was no water buffering changes in temperature, the stimulus was switched at shorter intervals and the probed temperature space was larger as well. This allowed for fewer samples overall to result in well-structured filters. Stimulus temperatures in *°*C were drawn from a Gaussian distribution:

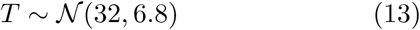

Temperature values were switched at random times with intervals in ms drawn from a gaussian distribution as well:

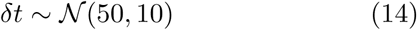

As during navigation simulations, the executed behaviors were used to derive the behavioral history input to the network during the simulations. For each network 10^7^ timesteps were simulated and the average temperatures in the 4 s preceding each turn or straight swim were computed.

#### Unit response clustering

All clustering was performed on the temperature branch of the networks. To cluster artificial neural network units into response types, the same temperature stimulus was presented to the networks that was previously used to analyze temperature representation in larval zebrafish and the same clustering approach was subsequently employed as well (Haesemeyer et al., 2018). Specifically, the pairwise correlations between all unit responses across all networks of a given type were calculated. Subsequently spectral clustering was performed with the correlation matrix as similarity matrix asking for 8 clusters as this number already resulted in some clusters with very weak responses, likely carrying noise. The cluster means were subsequently used as regressors and cells were assigned to the best-correlated cluster with a minimal correlation cutoff of 0.6. Cells that did not correlate with any cluster averages above threshold were not assigned to any cluster.

For *C. elegans* ANN units the same stimulus was used for clustering and temperature ramp responses were displayed for these obtained clusters.

To assign units in the mixed branch to clusters in the analysis of the retraining experiments, the same temperature stimulus was presented to the zebrafish ANN while speed and delta-heading inputs were clamped at 0. Correlation to cluster means of the temperature branch, again with a cut-off of 0.6, was subsequently used to assign these units to types.

#### Connectivity

Connectivity was analyzed between the first and second hidden layer of the temperature branch. Specifically, the average input weight of a clustered type in the first layer to a clustered type in the second layer was determined. The average weight as well as standard deviation across all networks and units was determined. If the standard deviation of a connection was larger than the average weight, the weight was set to 0.

#### Ablations and re-training of neural networks

Network ablations were performed by setting the activations of ablated units to 0 irrespective of their input. Retraining was performed using the same training data used to originally train the networks, and evaluating predictions using the same test data set. During retraining, activations of ablated units were kept at 0 and weight and bias updates of units were restricted to either the hidden layers in the temperature branch or in the mixed branch.

To identify unit types in the temperature or mixed branch after ablation or re-training, correlations to the corresponding cluster averages were used while presenting the same temperature stimulus used for clustering to the temperature branch and clamping the speed and delta-heading branch to 0.

#### Comparison of representations by PCA

To compare stimulus representations across all units and networks, our standard temperature stimulus was presented to all networks. Units from all networks and the types to compare (zebrafish thermal navigation vs. zebrafish phototaxis or zebrafish vs. *C. elegans* thermal navigation) were pooled and principal component analysis was performed across units. The first four principal components captured more than 95 % of the variance in all cases and were therefore used for comparison by evaluating the density along these principal components across network types.

#### Identification and mapping of zebrafish response types

Since the same temperature stimulus used for clustering network responses was previously presented to head embedded larval zebrafish (Haesemeyer et al., 2018) the cluster average responses were used as regressors to probe the larval zebrafish dataset. All neurons with a response correlated *>* 0.6 to these regressors was considered part of the response type (but we note that even when increasing or decreasing this threshold no neurons could be identified that matched the ON-OFF type).

To map these neurons back into the zebrafish brain we made use of the reference brain mapping generated in the imaging study.

### Code and data availability

The full source code of this project will be made available on github and all training data will be deposited upon final acceptance of the manuscript.

## Supplemental Figures

**Figure S1:**
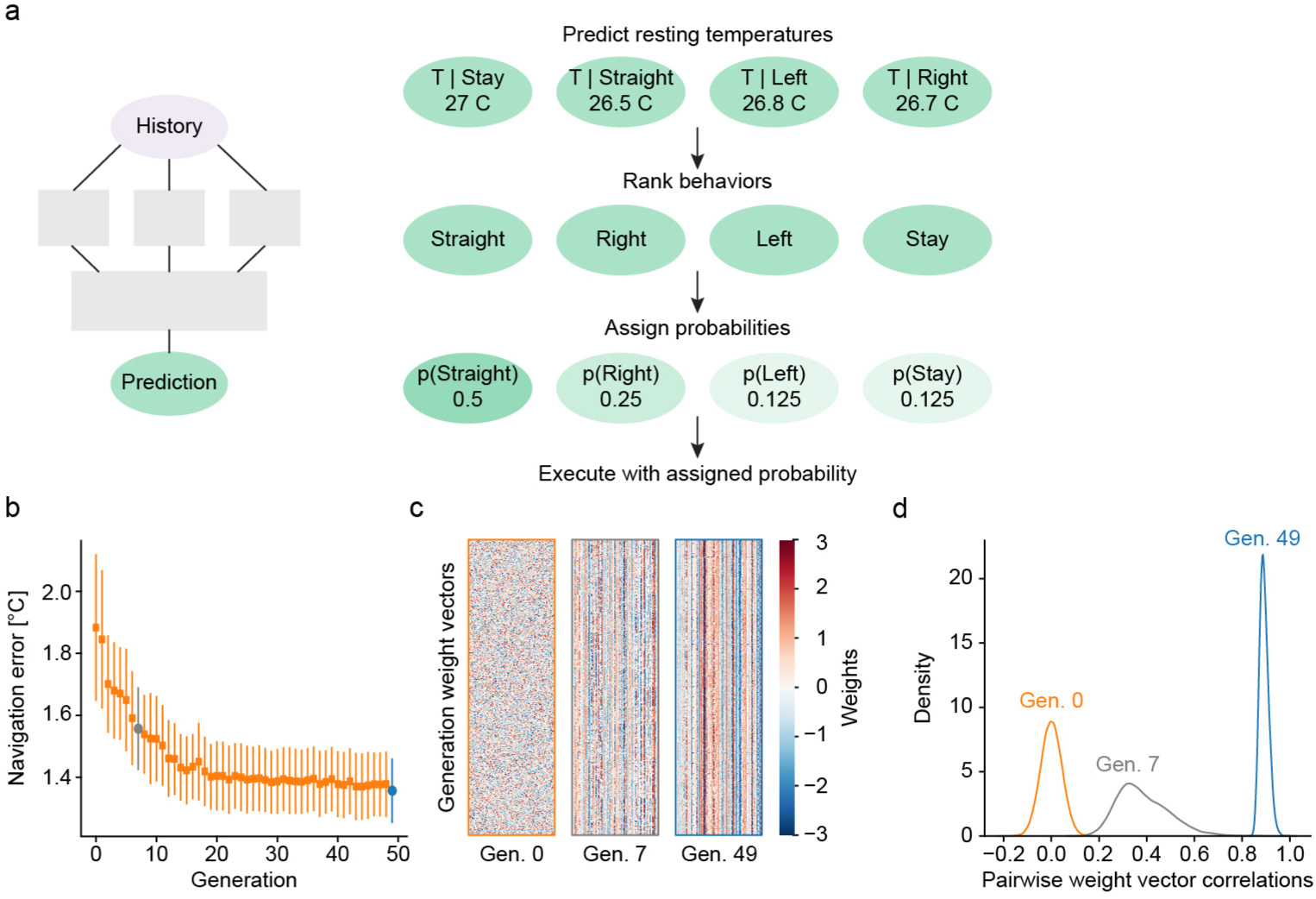
Network mechanics and evolution a) Schematic representation of how network predictions are transformed into behavioral selections during navigation. **b-d)** Example of p(Swim) weight evolution process on one network. b) Navigation error as average deviation from the desired temperature across all weight vectors in each generation. Error bars indicate standard deviation, generation 7 highlighted in grey and generation 49 in blue. c) Heatmap of all weight vectors in the indicated generations. Note the progressive increase in similarity. d) Pairwise correlations between all weight vectors in the indicated generations. Note the convergence on one solution during the evolutionary process.

**Figure S2:**
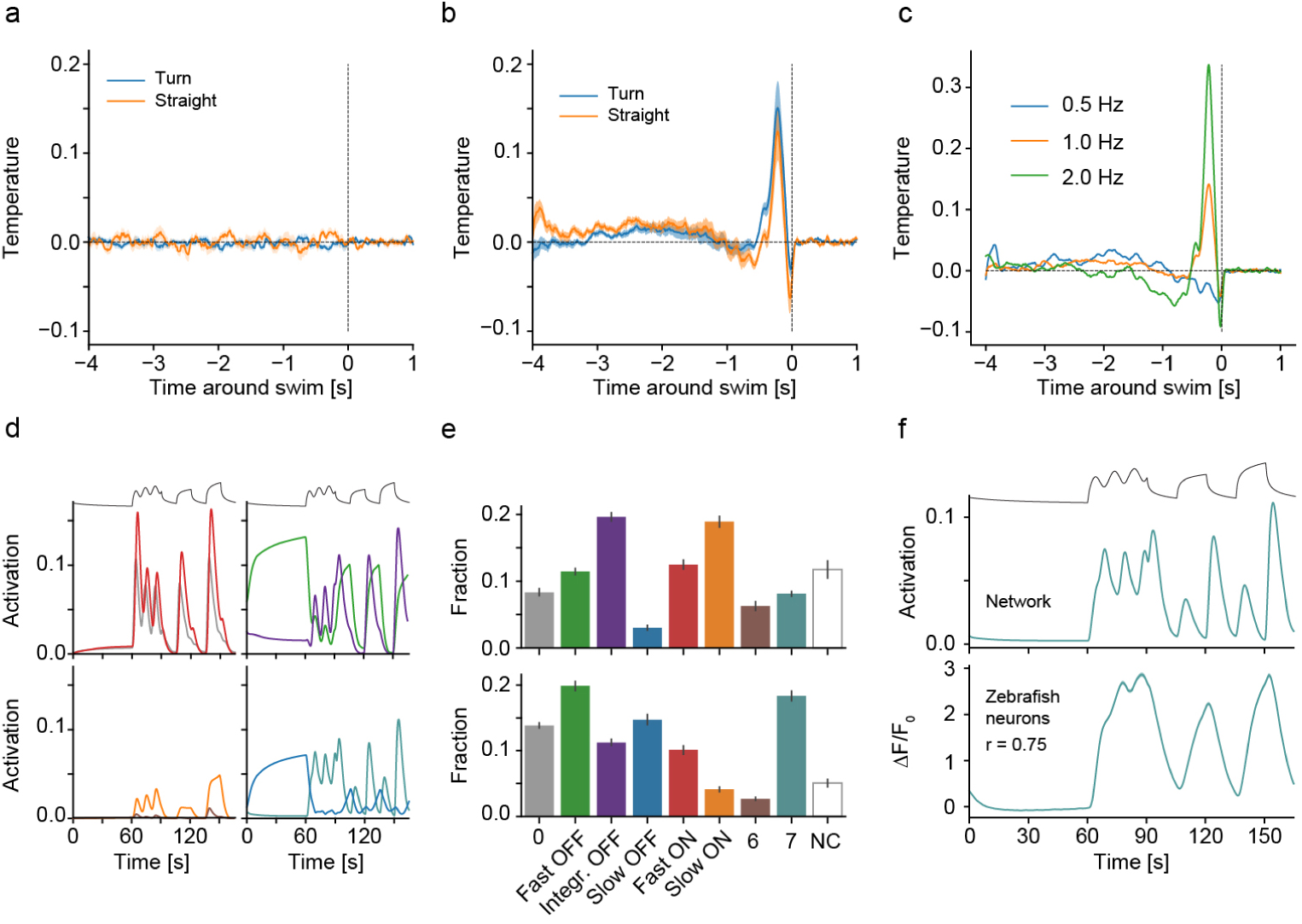
Characterization of zebrafish heat gradient navigation network a) Stimulus triggered averages for straight swims (orange) and turns (blue) when presenting white noise temperature stimuli to naïve networks. Dashed vertical line indicates time of swim. b) Stimulus triggered averages for straight swims (orange) and turns (blue) when presenting white noise temperature stimuli to fully trained and evolved networks. Dashed vertical line indicates time of swim. c) Stimulus triggered averages summed across all behaviors when training the models on training data with a baseline movement frequency of 0.5 Hz (blue), 1 Hz (orange) or 2 Hz (green). d) Average responses of all clusters to the heat stimulus depicted on top. Left two panels depict the four ON types present in the network, right two panels the four OFF types. Note that the brown ON type in the bottom left panel hardly response to the stimulus. e) Fraction of units contained in each cluster as well as fraction of units not belonging to any cluster (NC). f) The ON-OFF cell type present in the network, (teal) was used as a regressor (top panel) to identify the most closely matched cell types in zebrafish data (bottom panel, shading indicates bootstrap standard error across 1707 zebrafish neurons). Note that the identified zebrafish neurons do match network responses and in particular lack the OFF response present in the ON-OFF network type. Shading and error bars in all panels indicate bootstrap standard error across 20 networks.

**Figure S3:**
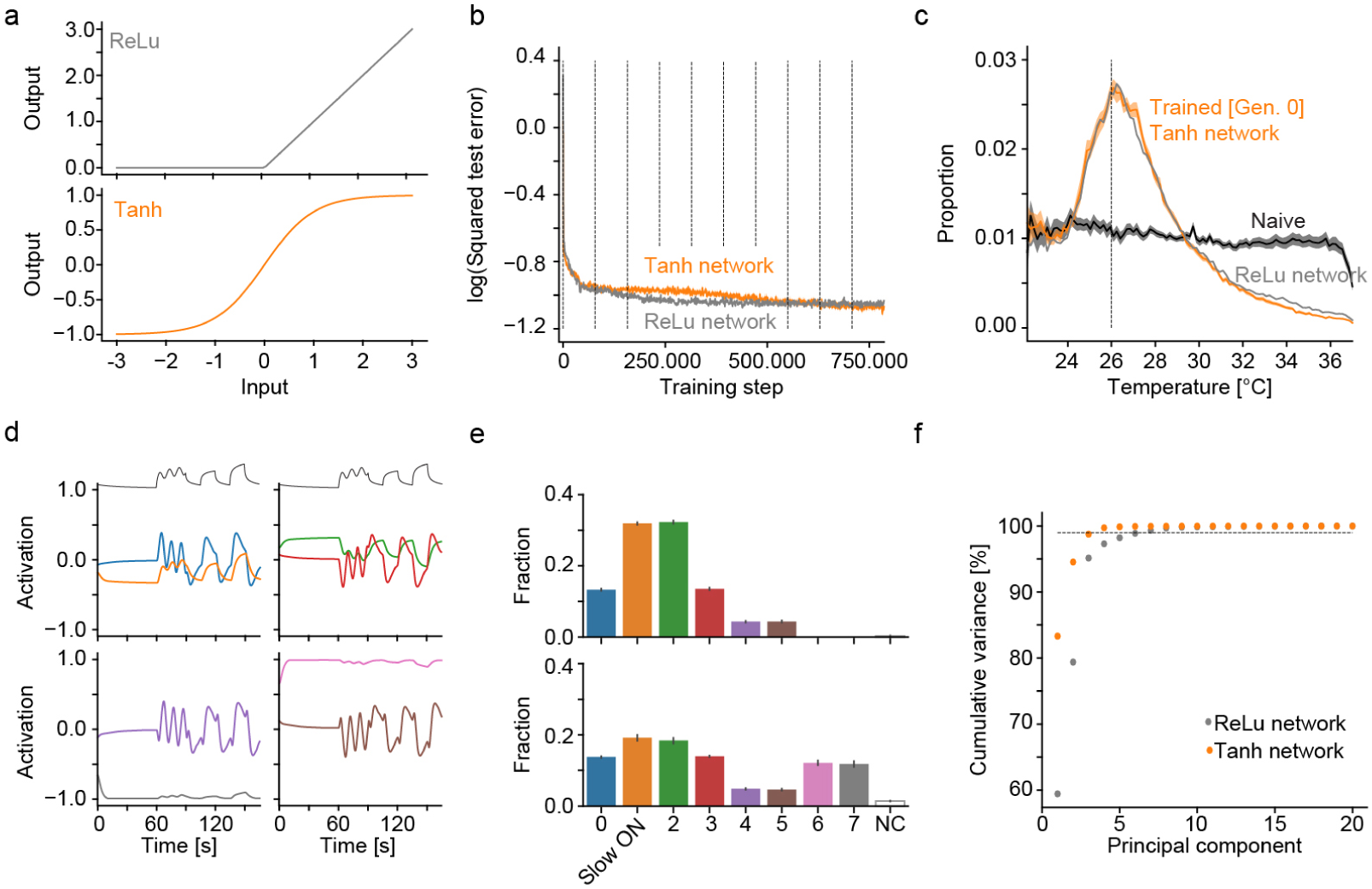
Influence of network activation function on temperature representation a) Top: Illustration of the ReLu activation function used in the networks of this paper except this figure. This activation function suppresses negative activations, similar to real neurons which cannot have negative firing rates. Bottom: Illustration of the Tanh activation function used for comparison, which is symmetric around 0, reporting both positive and negative responses. b) Log of the squared error in predictions on test data after the indicated number of training steps (dashed vertical lines demarcate training epochs) for the zebrafish heat navigation network with ReLu units (grey, replotted from Figure 1) and the same network with Tanh units. c) Occupancy in a radial heat gradient of naive Tanh (black), trained Tanh (orange) and trained Relu (grey, replotted from Figure 1) networks. Dashed vertical line at 26 *°*C indicates desired temperature. d) Average responses of all clusters in Tanh network to the heat stimulus depicted on top. Left two panels depict the four ON types present in the network, right two panels the four OFF types. Note that the OFF types in this case are just mirror-symmetric to the ON types. e) Fraction of units contained in each cluster as well as fraction of units not belonging to any cluster (NC). Error bars indicate bootstrap standard error across 20 networks. Note that the corresponding, mirror-symmetric OFF types are just as prevalent as their cognate ON types. f) Cumulative explained variance by principal components across all network units in 20 independently trained networks for ReLu units (grey) and Tanh units (orange). Dashed grey lines indicates 99 % of variance explained which is reached with 4 principal components in Tanh and 7 in ReLu networks. Shading and error bars in all panels indicate bootstrap standard error across 20 networks.

**Figure S4:**
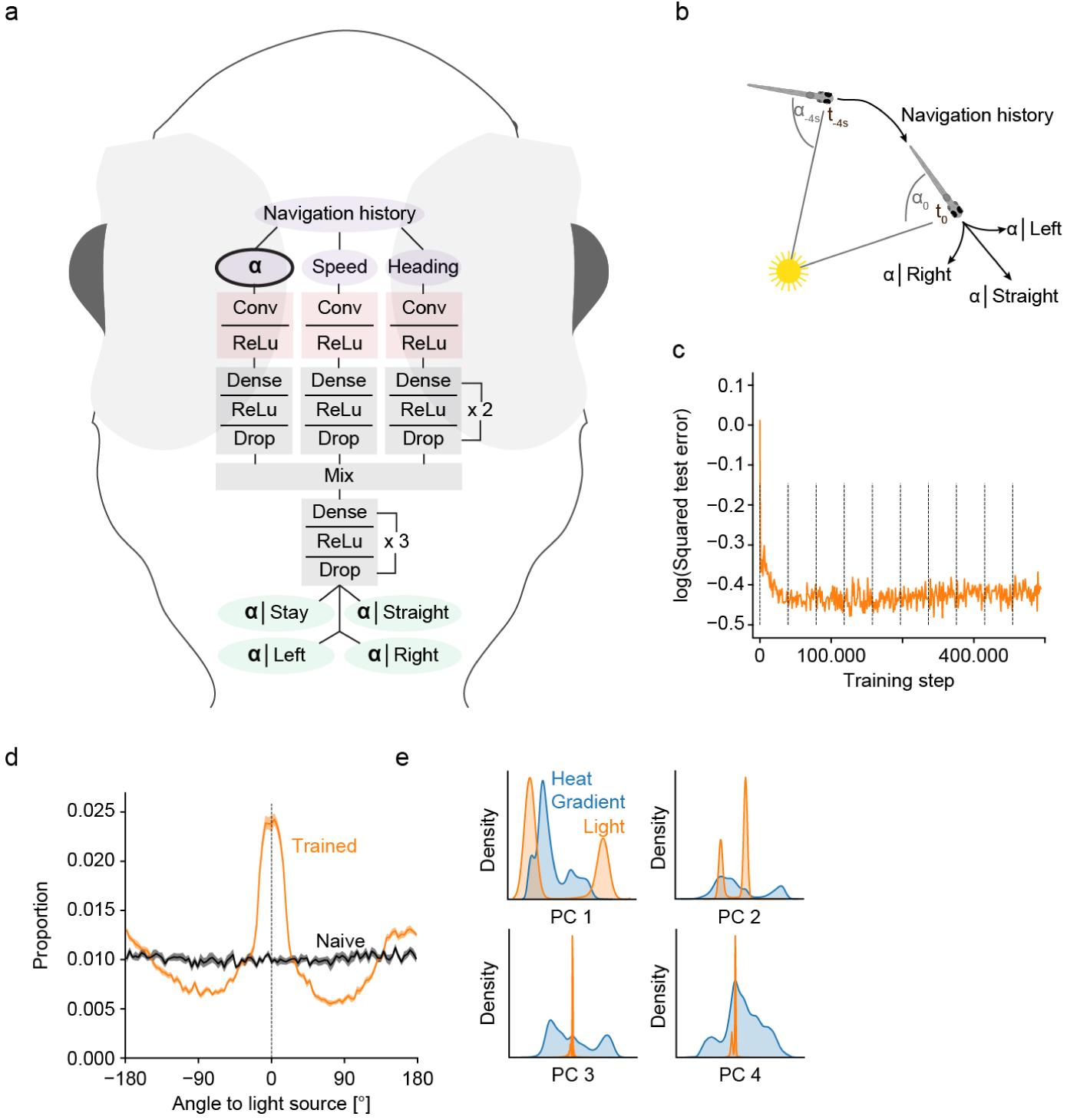
A network emulating zebrafish phototaxis a) Structure of the convolutional deep network for zebrafish phototaxis. Note that the architecture as exactly the same as in Figure 1d, only the input and prediction variables are angles instead of temperature. b) Schematic of the task of the phototaxis ANN: The network uses information about a 4s history of the relative angle to a light source and behavioral history to predict the angular position of the light source after different behavior selections. c) Log squared error of predictions on test data set during training. d) Performance when using the phototaxis network to favor behaviors that make a virtual fish face a light source (0 degree angular position) before (black line) and after training (orange line). Shading indicates bootstrap standard error across 14 networks. e) Comparison of all unit responses in the temperature branch of the zebrafish heat gradient ANN and the phototaxis ANN in PCA space when presenting the same time varying stimulus used in Figure 2b to both networks. The first four principal components capture *>* 95 % of the variance. Plots show occupational density along each PC for the gradient network (blue) and the phototaxis network (orange).

**Figure S5:**
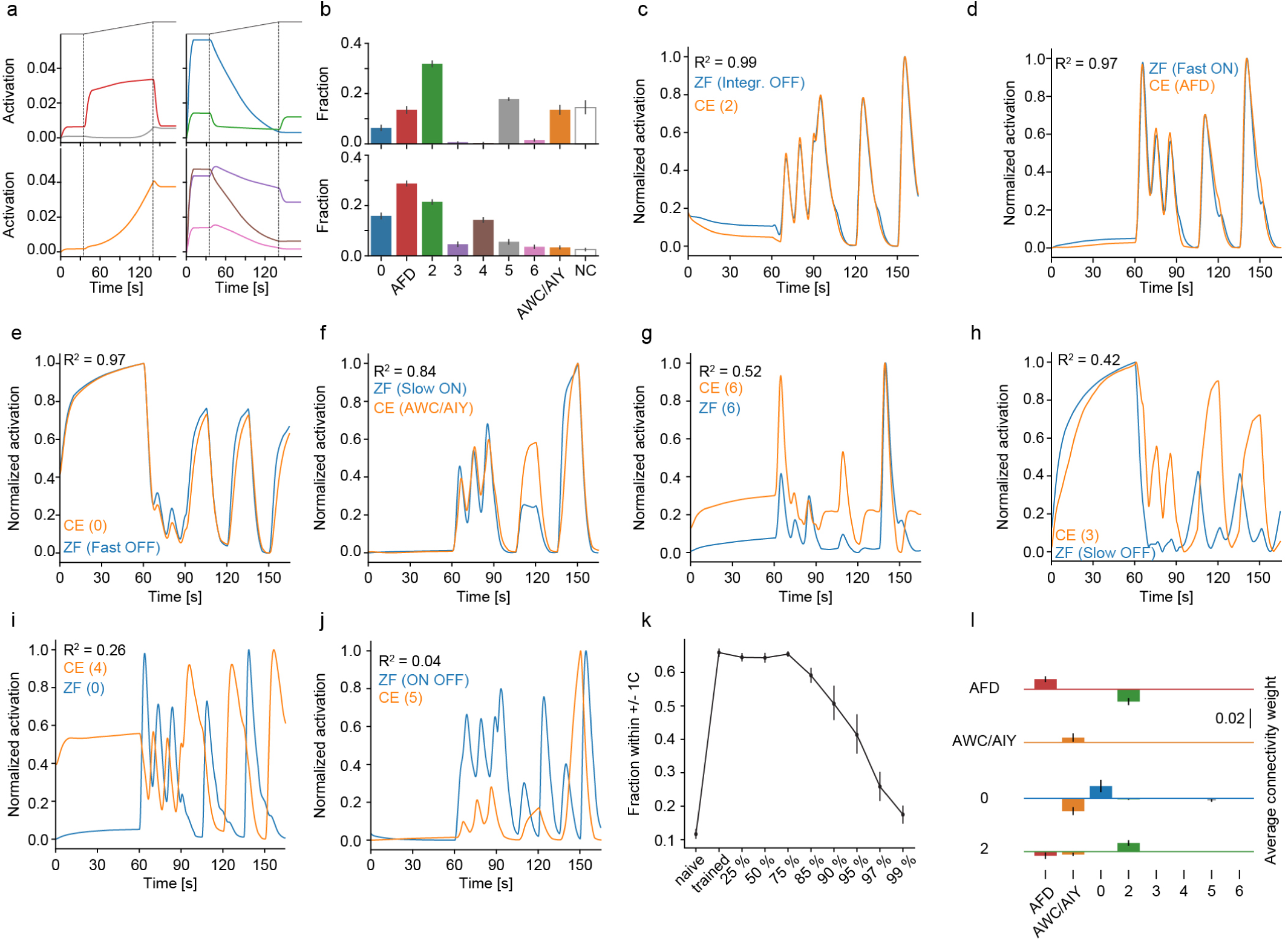
Characterization of *C. elegans* heat gradient navigation network a) Average responses of all clusters to the heat stimulus depicted on top. Left two panels depict the three ON types present in the network, right two panels the five OFF types. b) Fraction of units contained in each cluster as well as fraction of units not belonging to any cluster (NC). Error bars indicate bootstrap standard error across 20 networks. **c-j)** Comparison of zebrafish and *C. elegans* thermotaxis network response clusters. Each plot shows the average response of the indicated zebrafish network cluster (blue) and *C. elegans* network cluster (orange) with the coefficient of determination depicted on top. Pairs are ordered by decreasing correlation. k) Effect of random unit ablations on gradient navigation performance as fraction within 1 *°*C of desired temperature. Shown is performance for naive, fully trained and for random ablations of the indicated fraction of units in the temperature branch for *C. elegans* networks. Error bars indicate bootstrap standard error across 20 networks. l) Connectivity weights between layer 1 neuron types in the temperature branch (along x-axis) feeding into the indicated types of layer 2 neurons (panels). Types identified in Figure 4 are indicated by corresponding colored bars and the remaining four clusters are indicated by thinner gray bars on the right side. Error bars indicate standard deviation.

